# Mining the *Pseudomonas aeruginosa* genome for regulators of type III secretion system gene expression using FT-Tn-seq

**DOI:** 10.64898/2026.09.16.751967

**Authors:** Sardar Karash, Timothy L. Yahr

## Abstract

The *Pseudomonas aeruginosa* type III secretion system (T3SS) is an important virulence determinant used to mute and/or impair host immune defenses. Traditional transposon mutagenesis screens have been instrumental in identifying genes required for T3SS gene expression but our understanding of how those genes orchestrate regulatory control remains incomplete. A comprehensive inventory of the genes involved in the control of T3SS gene expression would help define regulatory mechanisms and provide additional targets for anti-virulence therapeutics. Here, we designed a fluorescently tagged transposon mutagenesis screen (FT-Tn-seq) and identified over 70 new genes important for T3SS gene expression. A role for 31 of those genes was validated using a secondary screen. Several candidate genes were selected for further studies. We demonstrate that deletion of *shaC*, which encodes a sodium-proton antiporter subunit, results in a significant defect in T3SS gene expression. That defect results from a combined effect on *exsA* transcription and ExsA translation. Deletion of *wspF*, a cyclic-di-GMP phosphodiesterase, results in a significant reduction of T3SS expression in strain PAK, but not in PA103. The effect on the T3SS is likely due to elevated levels of c-di-GMP in PAK. Finally, we identified a new sRNA (ivy) that participates in the control of the T3SS. Expression of ivy inhibits T3SS gene expression and requires the RNA-chaperone Hfq for inhibitory activity. Findings from this study will open new research avenues to better understand regulation of T3SS gene expression in *P. aeruginosa* and develop T3SS-specific therapeutic interventions.

**IMPORTANCE:** Anti-virulence therapeutic approaches are an alternative to conventional antibiotics. Rather than targeting an essential cellular pathway, anti-virulence approaches target a critical virulence function. The *P. aeruginosa* type III secretion system is a critical virulence determinant and validated target for anti-virulence approaches. In principle, any factor/pathway required for T3SS gene expression is a candidate target for therapeutic intervention. To advance our understanding of factors that control T3SS gene expression and broaden the list of potential therapeutic targets we performed a high-throughput screen to comprehensively identify factors required for T3SS gene expression. We now implicate 70 new genes in the control of T3SS gene expression.

## INTRODUCTION

*P. aeruginosa* is an opportunistic pathogen of humans, and a leading cause of nosocomial pneumonia, bloodstream, urinary tract, and post-operative infections (1, 2). One of the most important *P. aeruginosa* virulence determinants is a type III secretion-injectisome system (T3SS). The T3SS is required for full virulence in animal infection models and functions by translocating effector proteins with anti-phagocytic and cytotoxic properties into host cells (3–5). Mutants lacking a functional T3SS are significantly attenuated in pneumonia and burn wound infection models (6, 7).

The T3SS consists of ∼40 core genes that mostly encode for structural components of the secretion and translocation machinery but also include regulatory factors, the translocated effectors, and effector-specific chaperones (8). Expression of the T3SS regulon is controlled by ExsA, an activator of transcription (9, 10). Under non-inducing conditions, ExsA is complexed with ExsD in an inactive state and T3SS gene expression is low (11). Inducing conditions, such as growth in medium treated with a calcium chelator or physical contact of *P. aeruginosa* with host cells, result in secretion/translocation of protein substrates (12, 13). One of the secreted/translocated proteins is ExsE (14–16), a small protein that regulates ExsA-dependent transcription through a partner-switching mechanism (17). Partner-switching is an inherent regulatory mechanism of the T3SS that serves to couple secretory activity to gene expression and operates as follows. ExsE forms a complex with ExsC in the cytoplasm (18). Inducing conditions trigger secretion/translocation of ExsE. Removal of ExsE from the cytoplasm liberates ExsC and partner-switching ensues wherein ExsC forms a new interaction with ExsD, thereby disrupting the ExsD-ExsA complex (19). Release of ExsA from the ExsD-ExsA complex results in recruitment of ExsA to T3SS promoters and high levels of T3SS gene expression (19). Because T3SS gene expression is intimately coupled to secretory activity, mutations that disrupt secretory function result in ExsE accumulation in the cytoplasm and T3SS gene expression is low (15).

Partner-switching is an important mechanism for controlling T3SS gene expression but does operate in isolation. Several other regulatory systems contribute to control of T3SS gene expression by influencing *exsA* transcription or translation. Vfr is a cAMP-dependent transcription factor that stimulates *exsA* transcription through the P*_exsA_* promoter (20). P*_exsA_* promoter activity is also influenced by the small histone-like proteins MvaT/MvaU, the nucleoid associated protein Fis, and the quorum sensing regulator VqsM (21–23). ExsA translation is stimulated by the small RNA-binding protein RsmA and the RNA helicase DeaD, and is inhibited by the ribosomal protein L9 (RplI), Hfq in collaboration with small non-coding RNAs (sRNAs) 0161 and 179, and the heat shock protein CspC (24–29). In addition to direct control of *exsA* transcription or translation, many cellular products including tryptophan metabolites, spermidine, pyocins, and cyclic-di-GMP influence T3SS gene expression through mechanisms that remain to be fully delineated (17).

The core T3SS genes are encoded on a contiguous region of the genome and were identified using transposon mutagenesis screens (30–34). Many non-T3SS genes that contribute to T3SS gene expression were also identified by transposon mutagenesis screens (24, 33–40). While successful in identifying factors important for T3SS activity, those screens were non-comprehensive, time consuming, labor intensive, and biased towards genes with the strongest phenotypes. In the present study we performed a fluorescently-tagged transposon sequencing (FT-Tn-seq) screen to comprehensively identify genes and sRNAs that influence T3SS expression in *P*. *aeruginosa* strains PA103 and PAK. At a system-wide level we screened 152,584 unique insertion mutants in both strains and assessed the effect of those disruptions on T3SS gene expression. In addition to known T3SS genes, the screen identified up to 70 new genes and sRNAs that contribute to T3SS gene expression. We show that deletion of *shaC* inhibits T3SS expression by reducing *exsA* transcription and translation. Comparative genomics between the tested strains suggest that elevated c-di-GMP in the *wspF* mutant inhibits T3SS in the PAK strain but not in PA103. We also report that sRNA ivy functions as an Hfq-dependent inhibitor of T3SS expression.

## RESULTS

### Development and validation of FT-Tn-seq to identify genes required for T3SS gene expression

Fluorescently-tagged transposon insertion site sequencing (FT-Tn-seq) utilizes a fluorescent reporter for a regulatory system combined with random transposon mutagenesis, fluorescence-activated cell sorting (FACS), and next generation sequencing to identify factors important for expression of the reporter gene. Several studies have utilized the green-fluorescent protein (GFP) gene as a reporter in single cell analyses of *P. aeruginosa* T3SS gene expression (20, 41, 42). We chose the yellow-green fluorescent protein mNeonGreen (mNG) as the reporter because it is brighter than GFP (43). We first determined whether a single-copy reporter provides adequate fluorescence signal for FACS. The exoenzyme S promoter region (P*_exoS_*) was used to drive mNG expression and the resulting reporter (P*_exoS_*-mNG) was integrated into the chromosome at the neutral Tn7 site of *P. aeruginosa* strains PA103 and PAK (Fig. 1A). The P*_exoS_* promoter region is activated by ExsA and was previously validated as a reliable reporter of T3SS gene expression (11, 14). To examine P*_exoS_*-mNG reporter activity, cells were cultured under noninducing (− EGTA) and inducing (+ EGTA) conditions for T3SS gene expression and examined for fluorescence signal using flow cytometry. While 96% of the PA103 cells were induced for P*_exoS_*-mNG reporter activity (Fig. S1A vs B), only 20% of the PAK cells demonstrated high levels of fluorescence (Fig. S1D vs E). Previous studies using fluorescence-based reporters have also observed biphasic expression of the *P. aeruginosa* T3SS, wherein only a fraction of the cell population is induced for reporter activity (20, 41, 42). The biphasic pattern varies from strain-strain and is influenced by the growth medium and temperature (20, 41, 42).

**Figure 1.**
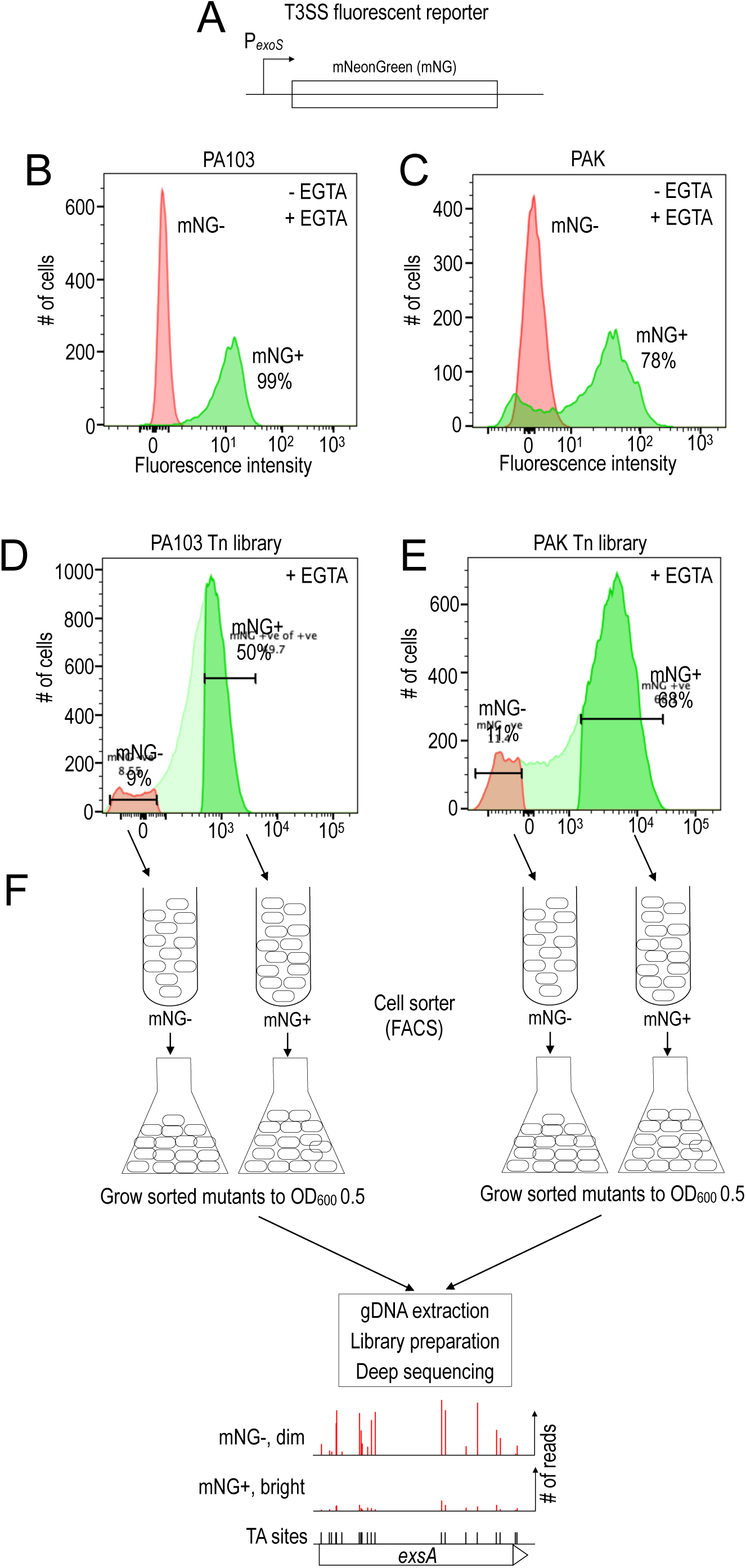
Outline of the FT-Tn-seq approach. **(A)** Diagram of the P*_exoS_*-mNeonGreen (P*_exoS_*-mNG) reporter used to sort dim (T3SS-Off) and bright (T3SS-On) cell populations. **(B-C)** Wild-type *P. aeruginosa* strains PA103 and PAK carrying the P*_exoS_*-mNG reporter were cultured separately under non-inducing (-EGTA, red) and inducing (+EGTA, green) conditions for T3SS gene expression. The percentage of cells scored as T3SS-On (mNG+) was determined using flow cytometry. **(D)** Transposon mutant libraries in strains PA103Δ*exoUT* and PAKΔ*exoSTY* were cultured under inducing conditions for T3SS gene expression and sorted into mNG-(T3SS-Off, red) and mNG+ (T3SS-On, green) populations. The sorted cells were cultured to an OD_600_ of 0.5, gDNA extracts were used for sequencing library preparation and deep sequencing. Fewer sequencing reads per gene in mNG-indicates the gene is required for T3SS expression.

The focus of the current study was to identify genes important for T3SS gene expression in response to low calcium (+ EGTA). To facilitate future studies examining T3SS gene expression in response to host cell contact, we constructed the transposon insertion libraries in PA103 and PAK mutant backgrounds lacking the primary T3SS effectors. Mutants lacking effectors (*exoU* and *exoT* for PA103 and *exoS, exoT,* and *exoY* for PAK) are essentially devoid of T3SS-dependent cytotoxicity towards cultured cells (44, 45). Deleting the effectors is necessary to prevent premature host cell lysis/death and to provide adequate time to induce detectable levels of T3SS gene expression. P*_exoS_*-mNG reporter activity was similar in the effectorless mutants when compared to their wt PA103 and PAK parents, demonstrating that T3SS gene expression is not affected by deleting the effectors (Fig. S1B vs C and E vs F). A single copy of the P*_exoS_*-mNG reporter, therefore, provides sufficient fluorescence signal to distinguish cells with low and high levels of T3SS gene expression.

The Mariner transposon used for this study generates high-density libraries by integrating at TA dinucleotide target sites (46). Transposon insertion libraries, generated in the PA103 Δ*exoUT* and PAK Δ*exoSTY* mutants carrying the P*_exoS_*-mNG reporter, were screened by FT-Tn-seq as outlined in Fig. 1. The PA103 Tn mutant library was cultured under inducing (2 mM EGTA) conditions for T3SS gene expression to OD_600_ 0.6. Growth to this density was sufficient to detect P*_exoS_*-mNG reporter activity in nearly 100% of the cells (Fig. 1 B, green). Separately, the same library was cultured under non-inducing (− EGTA) conditions to serve as input and gating controls for flow cytometry (Fig. 1B, red). The PA103 insertion mutants (52 million) were sorted by FACS and 9% of the dim cells (T3SS-Off, mNG-) and 50% of the bright cells (T3SS-On, mNG+) were collected (Fig. 1D and F). Sorting the PAK Tn library was more challenging than strain PA103 because the fluorescence of the P*_exoS_*-mNG reporter was not as strong in strain PAK and fewer cells were induced for T3SS gene expression owing to the stronger biphasic phenotype (Fig. S1E and F). To overcome this obstacle, we passaged the PAK Tn mutant library two times in the presence of 10 mM EGTA. This approach resulted in 78% of the cells being induced for P*_exoS_*-mNG reporter activity (Fig. S1G). The PAK mutants (66 million) were sorted, and 11% of the dim cells and 68% of the bright cells were collected for FACS (Fig. 1E and F).

The sorted dim and bright cells from both strains were grown separately to OD_600_ 0.5 and gDNA was extracted for Tn-seq library preparation (Fig. 1F). The PA103 Tn library had insertions in 82,787 of the 104,027 potential TA sites (80%) when sequenced to 750X coverage (Table S1). The PAK library had transposon insertions in 69,761 of the 95,792 potential TA sites (73%) when sequenced to 500X coverage (Table S1). To identify genes required for maximal T3SS gene expression, the fold change in gene reads was calculated by comparing the sorted dim mutants to the bright mutants. A negative Log_2_ fold change (Log_2_FC) indicates that the gene is required for maximal T3SS gene expression (Table S2 and S3).

### Genes with known roles in T3SS gene expression validate the FT-Tn-seq approach

As a first assessment of the data, we examined the T3SS genes. Because activation of the ExsECDA partner-switching mechanism and induction of T3SS gene expression requires a functional secretion system (17), we expected most of the T3SS genes to be overrepresented in the dim cell population. For strain PA103, 19 of the 36 T3SS genes demonstrated a Log_2_FC of at least −1.8 with a *P* value <0.05 (Table 1). Four of the T3SS genes (*pcr4, exsE, pscE, and pscI*) lacked TA sites and were not represented in the mutant library. Most (n = 9) of the remaining 13 genes also demonstrated negative Log_2_FC values but did not meet statistical measures of significance. For strain PAK only 8 of the T3SS genes demonstrated a Log_2_FC of at least −1.4 with a *P* value <0.05. Most of the remaining T3SS genes (n = 22), however, demonstrated negative Log_2_FC values but did not meet statistical measures of significance. We expected the data for strain PAK to be less robust than PA103 owing to the biphasic nature of T3SS gene expression in strain PAK, the need to passage the Tn-seq library under T3SS inducing conditions prior to FACS sorting, and the reduced sorting efficiency of the dim and bright cells relative to strain PA103. Each of those factors add noise/bias into the experimental workflow. Nevertheless, the finding that most (>92%) of the T3SS genes have negative Log_2_FC values demonstrates that the FT-Tn-seq approach can identify genes important for T3SS gene expression in strains PA103 and PAK. A few T3SS genes had positive Log_2_FC values. The T3SS gene with the largest positive value for both PA103 and PAK was *exsD*, consistent with its role as a negative regulator of T3SS gene expression (11).

**Table 1.** List of all genes in the type III secretion system regulon.

| <u>Gene Name</u> | <u>Strain</u><br><u>Locus Tag</u> | <u>PA103</u><br><u>Log<sub>2</sub>FC</u> | <u>PAK</u><br><u>Log<sub>2</sub>FC</u> | <u>Product name</u> |
| --- | --- | --- | --- | --- |
| *pscU* | PA1690 | **-2.86** | **-1.6** | secretion apparatus |
| *pscT* | PA1691 | **-3.51** | -1.57 | secretion apparatus |
| *pscS* | PA1692 | **-3.97** | -1.77 | secretion apparatus |
| *pscR* | PA1693 | **-3.24** | **-1.78** | secretion apparatus |
| *pscQ* | PA1694 | **-4.21** | -1.61 | secretion apparatus |
| *pscP* | PA1695 | **-3.59** | -2.2 | secretion apparatus |
| *pscO* | PA1696 | **-3.21** | -2.25 | secretion apparatus |
| *pscN* | PA1697 | **-2.6** | -1.68 | secretion apparatus |
| *popN* | PA1698 | **-2.61** | -1.04 | regulatory protein |
| *pcr1* | PA1699 | -1.17 | -1.57 | chaperone |
| *pcr2* | PA1700 | -2.52 | -1.79 | chaperone |
| *pcr3* | PA1701 | **-4.52** | -1.56 | chaperone |
| *pcr4* | PA1702 | * | * | chaperone |
| *pcrD* | PA1703 | **-3.53** | **-1.57** | secretion apparatus |
| *pcrR* | PA1704 | 0.32 | -1.04 | regulator |
| *pcrG* | PA1705 | 2.21 | -0.25 | regulator |
| *pcrV* | PA1706 | 0.02 | -0.72 | translocator |
| *prcH* | PA1707 | -0.09 | -0.85 | chaperone |
| *popB* | PA1708 | -0.83 | -1.26 | translocator |
| *popD* | PA1709 | -2.2 | -1.6 | translocator |
| *exsC* | PA1710 | -2.85 | -1.6 | partner-switching regulator |
| *exsE* | PA1711 | * | * | partner-switching regulator |
| *exsB* | PA1712 | **-3.93** | -1.38 | chaperone |
| *exsA* | PA1713 | **-3.27** | **-1.45** | partner-switching regulator |
| *exsD* | PA1714 | 0.82 | 2.64 | partner-switching regulator |
| *pscB* | PA1715 | -2.35 | 0.6 | chaperone |
| *pscC* | PA1716 | **-3.11** | **-1.56** | secretion apparatus |
| *pscD* | PA1717 | **-3.1** | **-1.74** | secretion apparatus |
| *pscE* | PA1718 | * | * | secretion apparatus |
| *pscF* | PA1719 | -3.11 | -1.96 | secretion apparatus |
| *pscG* | PA1720 | -3.33 | **-1.82** | secretion apparatus |
| *pscH* | PA1721 | **-1.83** | -1.82 | secretion apparatus |
| *pscl* | PA1722 | * | * | secretion apparatus |
| *pscJ* | PA1723 | **-3.63** | -1.66 | secretion apparatus |
| *pscK* | PA1724 | **-2.89** | -1.39 | secretion apparatus |
| *pscL* | PA1725 | **-2.94** | **-1.96** | secretion apparatus |

values in bold typeface have a p value <0.05
\*no TA present within the coding sequence of the indicated gene

We next examined non-T3SS-associated genes with known roles in the control of T3SS gene expression. The P*_exsA_* promoter is activated by Vfr, a cAMP-dependent transcription factor(20). cAMP is synthesized by the CyaB adenylate cyclase and CyaB activity influenced by the Pil/Chp chemosensory system (47). Once synthesized, cAMP stimulates Vfr-dependent transcription of *exsA*. As expected, *vfr*, *cyaB*, and 10 genes of the Pil/Chp system had negative Log_2_FC values (*P* < 0.05) in strain PA103 (Table 2). Most of the Pil/Chp genes identified in the PA103 dataset were previously implicated in the control of cAMP production (48). For strain PAK, *vfr* and *cyaB* were the only genes with negative Log_2_FC changes of at least −1.0 (*P* < 0.05). *tonB3* was previously implicated in extracellular pili assembly, and *gshAB* is also thought to play a role in Pil/Chp system regulation (49, 50). All three of those genes were identified in strain PA103.

**Table 2.** Non-T3SS genes that regulate T3SS gene expression.

| **cAMP/Vfr system** |  | **PA103** | **PAK** |  |
| --- | --- | --- | --- | --- |
| <u>Gene Name</u> | <u>Locus Tag</u> | <u>Log<sub>2</sub>FC</u> | <u>Log<sub>2</sub>FC</u> | <u>Product name</u> |
| *chpA* | PA0413 | **-1.88** | 0.12 | chemotactic signal transduction |
| *cyaB* | PA3217 | **-2.39** | **-1.1** | adenylate cyclase |
| *fimL* | PA1822 | **-2.5** | -1.1 | hypothetical protein |
| *fimV* | PA3115 | **-2.58** | **-0.2** | Motility protein FimV |
| *pilG* | PA0408 | -2.5 | -0.82 | type 4 fimbrial biogenesis |
| *pilH* | PA0409 | **-1.44** | -0.33 | type 4 fimbrial biogenesis |
| *pilI* | PA0410 | **-2.62** | -0.8 | type 4 fimbrial biogenesis |
| *pilJ* | PA0411 | **-2.2** | -0.43 | type 4 fimbrial biogenesis |
| *pilK* | PA0412 | **-2.28** | -0.6 | type 4 fimbrial biogenesis |
| *pilX* | PA4553 | **-1.28** | 0.52 | type 4 fimbrial biogenesis |
| *pilY1* | PA4554 | **-1.15** | 0.49 | type 4 fimbrial biogenesis protein |
| *vfr* | PA0652 | **-2.46** | **-1.35** | Cyclic AMP receptor-like protein |
| **influence *exsA* transcription or translation** |  |  |  |  |
| <u>Gene Name</u> | <u>Locus Tag</u> | <u>Log<sub>2</sub>FC</u> | <u>Log<sub>2</sub>FC</u> | <u>Product name</u> |
| *vqsM* | PA2227 | 1.25 | NP | transcriptional regulator VqsM |
| *fis* | PA4853 | **0** | -1.11 | DNA-binding protein Fis |
| *psrA* | PA3006 | 0.67 | 0.21 | transcriptional regulator PsrA |
| *deaD* | PA2840 | **-4.22** | -1.15 | ATP-dependent RNA helicase DeaD |
| *retS* | PA4856 | **-2.19** | **-1.06** | RetS regulator of Type III secretion |
| *rsmA* | PA0905 | 0.2 | 1.06 | RNA binding protein RsmA |
| *crc* | PA5332 | **-2.86** | **-1.52** | catabolite repression protein |
| **Spermidine transport** |  |  |  |  |
| <u>Gene Name</u> | <u>Locus Tag</u> | <u>Log<sub>2</sub>FC</u> | <u>Log<sub>2</sub>FC</u> | <u>Product name</u> |
| *spuE* | PA0301 | **-0.84** | -0.71 | polyamine transport protein |
| *spuF* | PA0302 | **-1.49** | **-0.82** | polyamine transport protein |
| *spuG* | PA0303 | **-1.48** | **-0.87** | polyamine transport protein |
| *spuH* | PA0304 | **-1.61** | **-0.85** | polyamine transport protein |
| **Cell envelope** |  |  |  |  |
| <u>Gene Name</u> | <u>Locus Tag</u> | <u>Log<sub>2</sub>FC</u> | <u>Log<sub>2</sub>FC</u> | <u>Product name</u> |
| *ctpA* | PA5134 | **-2.75** | -0.71 | C-terminal processing protease |
| *lbcA* | PA4667 | **-1.68** | -0.2 | lipoprotein binding partner of CtpA |
| *dsbA* | PA5489 | **-2.5** | -1.14 | thiol:disulfide interchange protein |
| **Undetermined role** |  |  |  |  |
| <u>Gene Name</u> | <u>Locus Tag</u> | <u>Log<sub>2</sub>FC</u> | <u>Log<sub>2</sub>FC</u> | <u>Product name</u> |
| *rne* | PA2976 | -2.95 | -1.45 | RNA metabolic process |
| *pchl* | PA4222 | -2.45 | -1 | component of ABC transporter |
| *pchH* | PA4223 | -1.12 | -1.26 | component of ABC transporter |
| *shaC* | PA1056 | -4.5 | -0.13 | sodium:proton antiporter activity |
| *gabD* | PA2065 | -1.44 | **-0.93** | oxidoreductase activity |
| **Genes with known roles that were not identified in the screen** |  |  |  |  |
| <u>Gene Name</u> | <u>Locus Tag</u> | <u>Log<sub>2</sub>FC</u> | <u>Log<sub>2</sub>FC</u> | <u>Product name</u> |
| *truA* | PA3114 | -0.94 | -0.46 | tRNA-pseudouridine synthase I |
| *ptrA* | PA2808 | -0.39 | -1.03 | Pseudomonas type III repressor A |
| *hfq* | PA4944 | essential | essential | regulation of RNA stability |
| *ptrB* | PA0612 | * | * | repressor, PtrB |
| *prtR* | PA0611 | essential | essential | transcriptional regulator PrtR |
| *tspR* | PA4857 | -0.28 | -0.09 | secretion by the type III |
| *nuoL* | PA2647 | -0.29 | -0.73 | NADH dehydrogenase I chain L |

values in bold typeface have a p value <0.05
\*no TA present within the coding sequence of the indicated gene

**Table 3.** New genes involved in T3SS gene expression.

| <u>Gene Name</u> | <u>Locus Tag</u> | <u>Log2FC</u> | <u>Product name</u> |
| --- | --- | --- | --- |
| *acpD* | PA3223 | -1.83 | FMN-dependent NADH-azoreductase |
| *apaH* | PA0590 | -1.54 | Bis(5'-nucleosyl)-tetraphosphatase, symmetrical |
| *arnF* | PA3558 | -1.38 | putative flippase subunit ArnF |
| *aruE* | PA0901 | -1.99 | Succinylglutamate desuccinylase |
| *cysQ* | PA5175 | -1.09 | 3'(2'),5'-bisphosphate nucleotidase CysQ |
| *erbR* | PA1978 | -1.53 | Transcriptional regulatory protein DegU |
| *fbpA* | PA5217 | -1.27 | putative binding protein component of ABC iron transporter |
| *gabD* | PA0265 | -1.44 | Glutarate-semialdehyde dehydrogenase |
| *galU* | PA2023 | -2.55 | UTP--glucose-1-phosphate uridylyltransferase |
| *gidA* | PA5565 | -2.37 | tRNA modification enzyme MnmG |
| *grxD* | PA3533 | -3.36 | Glutaredoxin 4 |
| *hel* | PA103-12375 | -1.4 | Lipoprotein e(P4) family 5'-nucleotidase |
| *hpcC* | PA4123 | -1.61 | NAD/NADP-dependent betaine aldehyde dehydrogenase |
| *hptB* | PA3345 | -1.88 | two-component system sensor histidine kinase HtpB |
| *hslU* | PA5054 | -1.53 | ATP-dependent protease ATPase subunit HslU |
| *htpG* | PA1596 | -1.64 | Chaperone protein HtpG |
| *kinB* | PA5484 | -1.7 | Alginate biosynthesis sensor protein KinB |
| *liuB* | PA2014 | -1.13 | Methylmalonyl-CoA carboxyltransferase 12S subunit |
| *metF* | PA0430 | -1.03 | 5,10-methylenetetrahydrofolate reductase |
| *mexA* | PA0425 | -1.53 | Multidrug resistance protein MexA |
| *mexB* | PA0426 | -1.5 | Multidrug resistance protein MexB |
| *narJ* | PA3873 | -1.08 | putative cofactor assembly chaperone NarW |
| *nosZ* | PA3392 | -1.02 | Nitrous-oxide reductase |
| *nuoD* | PA2639 | -1.73 | NADH-quinone oxidoreductase subunit C/D |
| *oadA* | PA5435 | -2.7 | Oxaloacetate decarboxylase alpha chain |
| PA0371 | PA0371 | -1.16 | putative Zn-dependent peptidase, M16 family |
| PA0372 | PA0372 | -1.32 | putative zinc protease |
| PA0429 | PA0429 | -1.03 | Glycosyl transferase |
| PA0431 | PA0431 | -1.95 | (p)ppGpp synthase/hydrolase, HD superfamily |
| PA0596 | PA0596 | -1.21 | N-acetylmuramate/N-acetylglucosamine kinase |
| PA0769 | PA0769 | -1.62 | DUF4845 domain-containing protein |
| PA0943 | PA0943 | -1.39 | Dehydrogenase |
| PA1011 | PA1011 | -1.11 | Outer membrane protein assembly factor BamC |
| PA103-01262 | PA103-05485 | -1.42 | hypothetical protein |
| PA1064 | PA1064 | -1.1 | Uncharacterized membrane protein |
| PA1140 | PA1140 | -1.25 | (S)-ureidoglycine aminohydrolase |
| PA1612 | PA1612 | -1.27 | ABC-type uncharacterized transport system |
| PA1766 | PA1766 | -1.975 | Glutathione synthase |
| PA1767 | PA1767 | -1.72 | Inactive transglutaminase |
| PA2108 | PA2108 | -1.17 | Putative thiamine pyrophosphate-containing protein YdaP |
| PA2111 | PA2111 | -2.42 | 5-oxoprolinase subunit B |
| PA2112 | PA2112 | -2.73 | 5-oxoprolinase subunit A 3 |
| PA2550 | PA2550 | -3.41 | (R)-benzylsuccinyl-CoA dehydrogenase |
| PA3052 | PA3052 | -1.77 | T2SSE-N domain-containing protein |
| PA3699 | PA3699 | -1.93 | DNA-binding transcriptional regulator YbjK |
| PA3894 | PA3894 | -1.48 | Toluene efflux pump outer membrane protein Ttgi |
| PA3978 | PA3978 | -1.46 | TPR repeat |
| PA4029 | PA4029 | -1.84 | Protein DedA |
| PA4423 | PA4423 | -1 | Penicillin-binding protein activator LpoA |
| PA4583 | PA4583 | -1.97 | RNA-splicing ligase RtcB |
| PA4629 | PA4629 | -1.89 | Putative Ca<sup>2+</sup>/H<sup>+</sup> antiporter, TMEM165/GDT1 family |
| PA4972 | PA4972 | -1.6 | DUF3298 domain-containing protein |
| PA5279 | PA5279 | -1.11 | DUF484 domain-containing protein |
| PA5436 | PA5436 | -1.14 | acetyl-CoA carboxylase biotin carboxylase subunit |
| PA5547 | PA5547 | -1.7 | Phosphoserine phosphatase |
| *pgi* | PA4732 | -2.32 | Glucose-6-phosphate isomerase |
| *pncA* | PA4918 | -1.35 | Nicotinamidase/pyrazinamidase |
| *ppiD* | PA1805 | -2.21 | Peptidyl-prolyl cis-trans isomerase D |
| *prfC* | PA3903 | -1.7 | Peptide chain release factor RF3 |
| *ptsP* | PA0337 | -1.4 | Phosphoenolpyruvate-dependent phosphotransferase |
| *pvdS* | PA2426 | -1.34 | pyoverdine signaling pathway sigma factor PvdS |
| *rbsA* | PA1947 | -2.87 | Arabinose import ATP-binding protein AraG |
| *rbsC* | PA1948 | -1.49 | Ribose import permease protein RbsC |
| *rlmN* | PA3806 | -1.08 | Dual-specificity RNA methyltransferase RlmN |
| *rluD* | PA4544 | -1.7 | Ribosomal large subunit pseudouridine synthase D |
| *rnd* | PA1294 | -1.1 | Ribonuclease D |
| *saiB* | PA0171 | -2.21 | biofilm formation regulator kinase SiaB |
| *shaA* | PA1054 | -3.37 | Na(+)/H(+) antiporter subunit A |
| *shaC* | PA1056 | -4.5 | Na(+)/H(+) antiporter subunit D |
| *smpB* | PA4768 | -1.81 | SsrA-binding protein |
| *ssrA* | ssrA | -1.54 | transfer-messenger RNA, SsrA |
| *sthA* | PA2991 | -2.49 | Si-specific NAD(P)(+) transhydrogenase |
| *tonB3* | PA0406 | -2.27 | Periplasmic protein TonB |
| *ygdP* | PA0336 | -1.25 | RNA pyrophosphohydrolase |

Control of ExsA synthesis is also an important regulatory checkpoint. The DeaD RNA helicase directly stimulates ExsA translation (24) and had a Log_2_FC of −4.22. RsmA is also thought to stimulate ExsA translation and is critical for T3SS gene expression (17). It was thus surprising that the Log_2_FC for *rsmA* was only 0.2, especially since *retS* was identified by the screen (Log_2_FC was −2.19). RetS indirectly influences RsmA activity by inhibiting *rsmYZ* transcription (51). RsmYZ are small non-coding RNAs that sequester RsmA from target RNAs expression (17). Thus, in the absence of *retS*, RsmYZ levels are elevated and RsmA availability is very low. A *rsmA* mutant, therefore, should phenocopy a *retS* mutant. The failure of the screen to identify *rsmA* likely reflects low coverage in the Tn library (only 1 of 3 the TA sites contained insertions) and the slow growth phenotype associated with loss of RsmA function.

While DeaD and RsmA play direct roles in *exsA* transcription or ExsA synthesis, many other genes influence T3SS gene expression through mechanisms that remain to be defined. Those genes include an ABC transporter (*pchHI*) required for virulence in *Dictyostelium* and for efficient expression of T3SS (39); a thiol:disulfide oxidoreductase (*dsbA*) essential for T3SS expression and intracellular survival in HeLa cells (52); a periplasmic protease (*ctpA*) necessary for T3SS function, and virulence in a mouse pneumonia model (53); and a sodium-proton antiporter (*shaA-F*), where deletion of *shaA* leads to reduced virulence in mice (24, 54). Each of the genes highlighted above were identified in the PA103 screen with Log_2_FC values of at least −1.12, though none met the *p* <0.05 threshold test for significance (Table 2).

### New genes implicated in the control of T3SS gene expression

To further validate the experimental data, all PA103 genes with a negative Log_2_FC value and *P* values ≤ 0.05 were compiled into a list of 146 candidates. Genes (N = 31) were randomly selected from the list and tested in a secondary screen by introducing the P*_exsD_-LacZ* reporter into transposon insertion mutants obtained from a *P. aeruginosa* strain PA14 mutant library (55). Similar to the P*_exoS_*-mNG reporter, P*_exsD_-LacZ* reporter activity is ExsA-dependent and exhibits robust induction under inducing conditions for T3SS gene expression (11). With only one exception (the *aruE* mutant), P*_exsD_-LacZ* reporter activity in each of the remaining mutant backgrounds demonstrated a significant reduction when compared to the parental strain (Fig. 2). Based on these findings we selected a Log_2_FC value ≤ −1.0 as a cutoff for candidate genes required for T3SS gene expression, which narrowed the list to 115 genes in strain PA103 (Table S2). Of these 115 genes, 73 represent new candidate genes for involvement in control of T3SS gene expression (Table 2). The remainder of the study focused on a few of the new genes.

**Figure 2.**
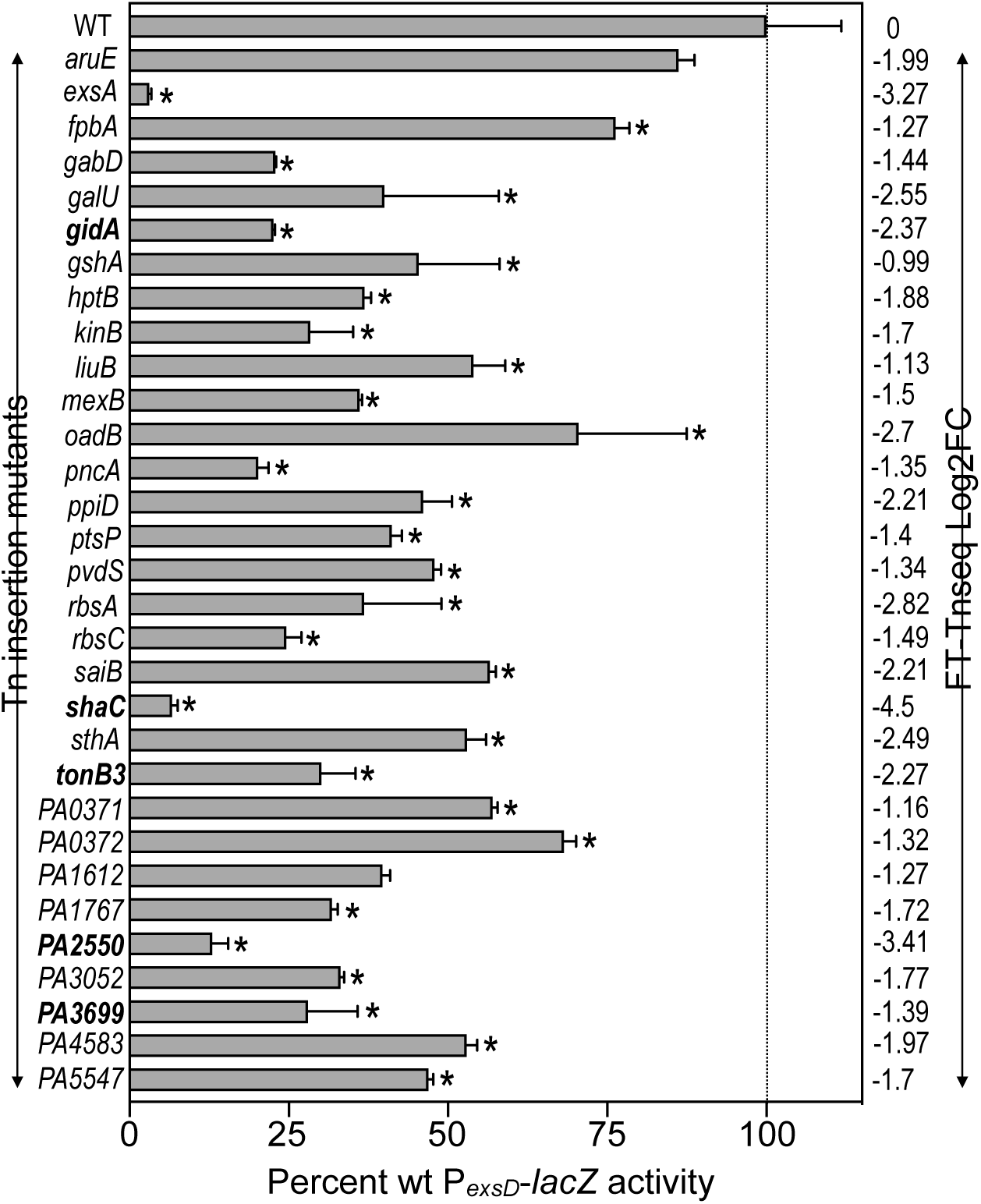
Validation of FT-Tn-seq with P*_exsD_-lacZ* reporter and RNA-seq. Secondary screen used to validate T3SS-related genes identified in the primary FT-Tn-seq screen. The P*_exsD_*-lacZ transcriptional reporter for T3SS expression was integrated into the indicated *P. aeruginosa* Tn mutants (N=30). Mutant and wt strains were grown under T3SS-inducing (+EGTA) conditions and assayed for P*_exsD_-lacZ* reporter activity. Beta-galactosidase activity was measured in Miller units with the standard error representing the average of at least three experiments. *, *P* < 0.05. The reported values are percent activity relative to wt cells carrying the P*_exsD_-lacZ* reporter.

### The sodium-proton antiporter (Sha) is required for T3SS expression

*P*. *aeruginosa* has four sodium-proton antiporters, thought to play a role in maintaining Na^+^ and H^+^ homeostasis (56). The Sha sodium-proton antiporter consists of 6 subunits (*shaABCDEF*). Two prior transposon mutagenesis screens identified *shaC* as being critical for maximal T3SS gene expression (24, 40). In our screen each of the *sha* genes, with the exception of *shaB*, demonstrated a negative Log_2_FC change, though only *shaA* and *shaC* met the threshold for significance (Log_2_FC ≤ −1.0). The transposon insertion profile implies that the entire *shaABCDEF* operon is required for T3SS gene expression. We constructed an in-frame *shaC* deletion mutant (Δ*shaC*) in the PA103 and PAK parental backgrounds and integrated a P*_exsD_-lacZ* transcriptional reporter at the ΦCTX phage attachment site. As expected, P*_exsD_-lacZ* reporter activity was significantly reduced in the PA103 and PAK Δ*shaC* mutants (Fig. 3A). Since both strains showed strong phenotypes, we only conducted follow-up experiments in strain PA103. Complementation of the Δ*shaC* mutant restored P*_exsD_-lacZ* reporter activity, demonstrating that the *shaC* deletion is responsible for the loss of T3SS activity (Fig. 3B). We also measured expression of the Vfr-dependent P*_exsA_-lacZ* reporter activity (20) and observed a significant reduction in the Δ*shaC* mutant. This finding was confirmed by immunoblotting for the ExsA protein (Fig. 3C). Finally, we integrated *exsA* under the control of a rhamnose-inducible promoter at the Tn7 site in Δ*exsA*-Δ*shaC* and Δ*exsA* mutants. While the inducer restored T3SS activity in the Δ*exsA* mutant, it did not restore T3SS expression in the Δ*exsA*-Δ*shaC* background (Fig. 1D). These results suggest that the T3SS defect in the Δ*shaC* background may occur on both the transcriptional level through the Vfr-dependent P*_exsA_* promoter and at the level of ExsA translation.

**Figure 3.**
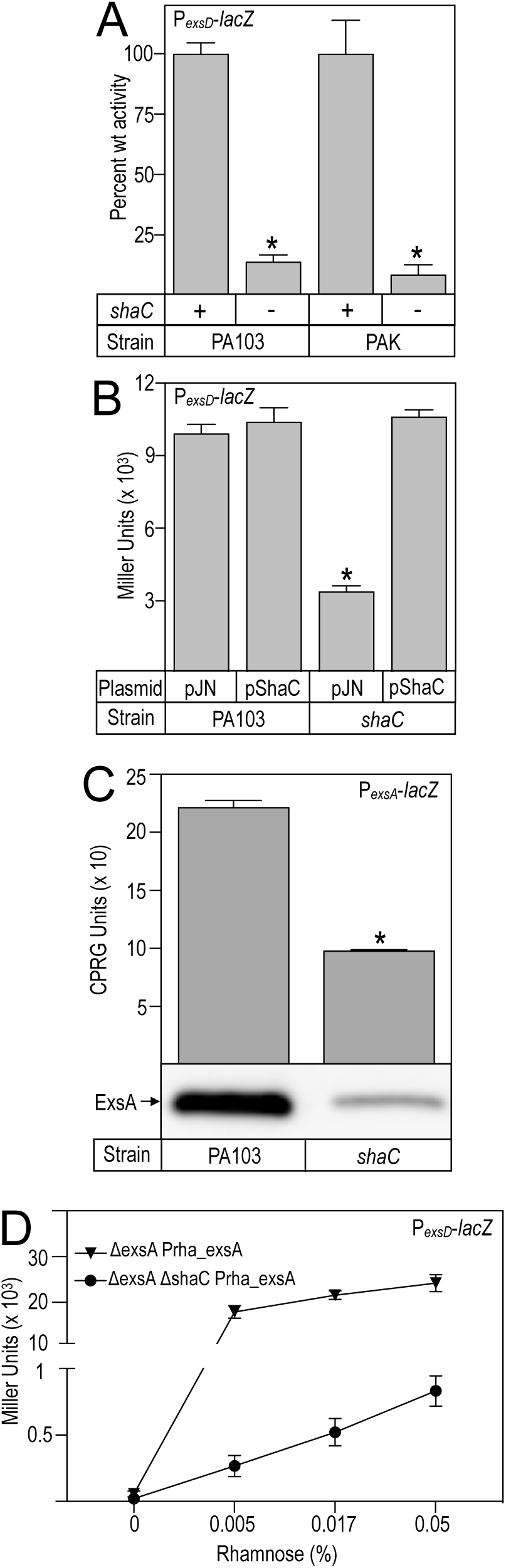
*shaC* is required for T3SS expression in *P. aeruginosa*. **(A)** Maximal P*_exsD_-lacZ* reporter activity is *shaC*-dependent in strains PA103 and PAK. The P*_exsD_-lacZ* reporter was inserted into wildtype and *ΔshaC* mutants to measure T3SS gene expression. The resulting strains were cultured under T3SS-inducing conditions (+EGTA) and assayed for P*_exsD_-lacZ* reporter activity. **(B)** WT strain PA103 and a *shaC* mutant carrying a P*_exsD_-lacZ* reporter were transformed with either a vector control (pJN105) or a *shaC* expression vector (pShaC). Strains were cultured under T3SS-inducing conditions and assayed for P*_exsD_-lacZ* reporter activity. **(C)** PA103 WT and *ΔshaC* carrying a P*_exsA_-lacZ* reporter were cultured under T3SS-inducing conditions and assayed for P_exsA_-*lacZ* reporter activity. Cell lysate fractions from the same cultures were immunoblotted for ExsA. **(D)** An *exsA* mutant and an *exsA shaC* double mutant carrying a rhamnose-inducible copy of *exsA* integrated at the Tn7 site were cultured under T3SS-inducing conditions. Rhamnose was included in the growth medium to induce *exsA* transcription, and T3SS expression was measured using the P*_exsD_-lacZ* reporter. For all experiments, the standard error represents the average of at least three biological replicates. * denotes *P* < 0.05.

### Inhibition of T3SS gene expression by elevated c-di-GMP in a *wspF* mutant is strain dependent

Cyclic-di-GMP (c-di-GMP) is a signaling molecule that influences many bacterial traits including biofilm formation and virulence. c-di-GMP is synthesized from two molecules of GTP by diguanylate cyclases (DGC) and is degraded by phosphodiesterases (PDEs). *P. aeruginosa* has ∼30 enzymes with DGC activity and ∼20 enzymes with PDE activity (57). As many as 5 DGC’s and 9 PDE’s may influence T3SS gene expression (17). The environmental signals and manner by which these enzymes coordinate c-di-GMP homeostasis remains poorly understood. One exception is the Wsp system, which stimulates c-di-GMP production through the WspR PDE in response to surface sensing (58). WspR activity is under negative feedback control by WspF. Cells lacking functional WspF, therefore, have increased levels of c-di-GMP and low T3SS gene expression would be expected. In our FT-Tn-seq screen, *wspF* was required for robust T3SS expression in the PAK background. The same was not true in the PA103 background as Tn insertions in *wspF* had no effect on T3SS gene expression. The latter finding implies that the *wspF* requirement is strain-specific.

We examined this further by deleting *wspF* in both strains and then measured P*_exsD_-LacZ* reporter activity. Consistent with FT-Tn-seq results, P*_exsD_-LacZ* reporter activity was significantly reduced in the PAK Δ*wspF* mutant and unaffected in the PA103 Δ*wspF* deletion mutant when compared to the parental strains (Fig. 4A). To verify that *wspF* deletion results in elevated levels of c-di-GMP production, a P*_cdrA_*-GFP reporter was introduced into the PAK and PA103 Δ*wspF* backgrounds (58, 59). P*_cdrA_*-GFP is a transcriptional reporter responsive to elevated c-di-GMP (60). P*_cdrA_*-GFP reporter activity was significantly elevated in the PAK Δ*wspF* mutant (Fig. 4B),but was unaffected in PA103 Δ*wspF* (data not shown). As alternative approach to increase c-di-GMP levels in strain PA103 we replaced the native *wspR* allele with a *wspR* D16N mutant allele. The D16N substitution results in constitutive WspR diguanylate cyclase activity. When c-di-GMP levels were tested using the P*_cdrA_*-GFP reporter, however, the *wspR* D16N had no effect on reporter activity (data not shown). It is unclear why the two engineered mutants (Δ*wspF* and *wspR* D16N) in strain PA103 had no effect on P*_cdrA_*-GFP reporter activity. While some studies find that elevated levels of c-di-GMP are associated with reduced T3SS activity (57, 61), at least one report found no relationship between c-di-GMP and T3SS gene expression (62). The findings from our screen add to this ambiguity.

**Figure 4.**
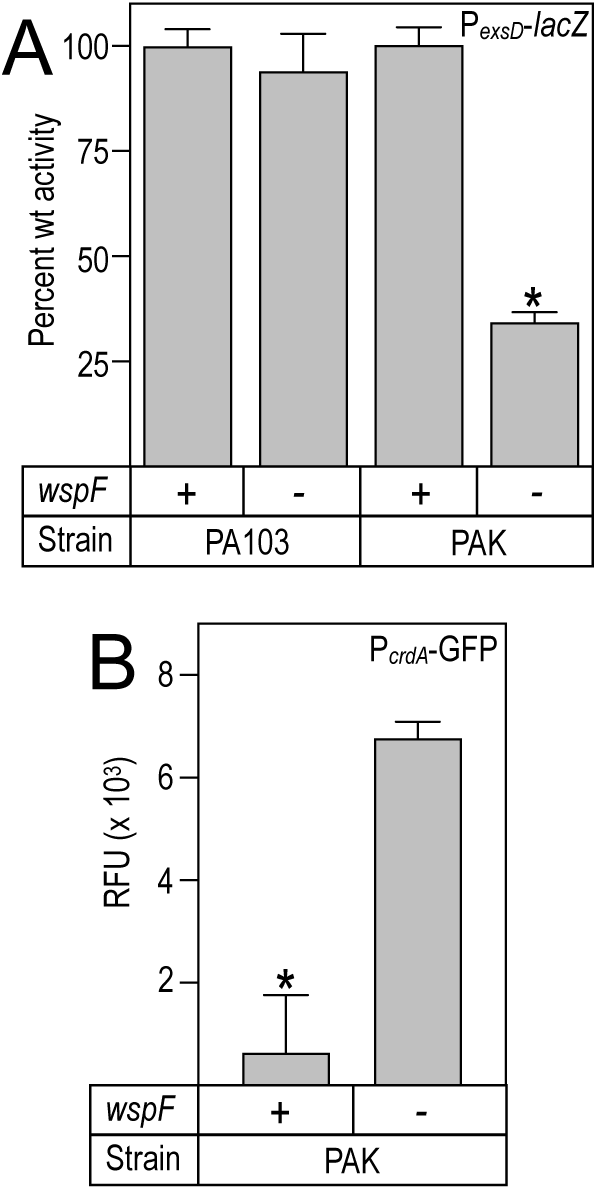
Reduced T3SS gene expression in the *wspF* mutant is strain specific. **(A)** The P*_exsD_-lacZ* reporter was inserted into to the indicated wildtype and Δ*wspF* mutants. Strains were cultured under T3SS-inducing conditions (+EGTA) and assayed for P*_exsD_-lacZ* reporter activity. **(B)** WT PAK and a Δ*wspF* mutant were transformed with a P*_crdA_*-GFP reporter transformed into to indirectly measure c-di-GMP levels. Strains were cultured under T3SS-inducing conditions, and GFP fluorescence was measured and reported as relative fluorescence units (RFU).

### sRNA ivy, a new Hfq-dependent inhibitor of T3SS gene expression

Small noncoding RNAs (sRNAs) play a critical role in regulation of T3SS gene expression (28). We combined the FT-Tn-seq results with several bioinformatics tools to mine for sRNAs that influence T3SS gene expression. First the Infernal pipeline (63) was used to search for sRNAs in the PA103 genome using the Rfam database resulting in the identification of 87 candidate sRNAs. Next, the genomic junctions obtained from the FT-Tn-seq dim and bright mutants were mapped to the 87 candidate sRNAs. From that pool 48 sRNAs had increased Log_2_FC values while 16 sRNAs had a reduced Log_2_FC. Of the remaining 23 sRNAs, 12 lacked TA insertion sites and 11 either did not tolerate Tn insertions or our sequencing did not cover the short sRNA sequence. The sRNAs were rank-ordered based on Log_2_FC values from highest to lowest (Table S4). The known sRNAs *rsmY* and *rsmZ* ranked 10 and 23 in the list, respectively. This is consistent with previous findings showing that mutants lacking either *rsmY* or *rsmZ* have decreased levels of T3SS gene expression (owing to increased RsmA availability) (28, 64) and served to validate our approach to identifying sRNAs that control T3SS gene expression.

To further validate whether the sRNA candidates influence T3SS gene expression, the 5 top candidates were cloned into an expression vector and introduced into wt *P. aeruginosa* PA103 carrying the P*_exsD_-LacZ* reporter. One of the candidates, sRNA ivy (65), demonstrated significant inhibition of P*_exsD_-LacZ* reporter activity (Fig. 5A). sRNA Ivy (ivy-DE, inhibitor of vertebrate lysozyme downstream element) is a 92-nucleotide RNA that was discovered in a previous bioinformatic analysis (65) and is only present in the *Pseudomonads*. Expression of ivy in strains PAK and PA103 results in strong inhibition of P*_exsD_-LacZ* reporter activity (Fig. 5B).The base pairing of sRNAs with target mRNAs is usually facilitated by RNA chaperones. In *P.aeruginosa* the primary RNA chaperone is Hfq. When expressed in a PA103 mutant lacking functional Hfq, the inhibitory activity of ivy was largely suppressed (Fig. 5C). The dependence of ivy on Hfq findings suggests that ivy inhibits T3SS gene expression by base-pairing with an RNA target.

**Figure 5.**
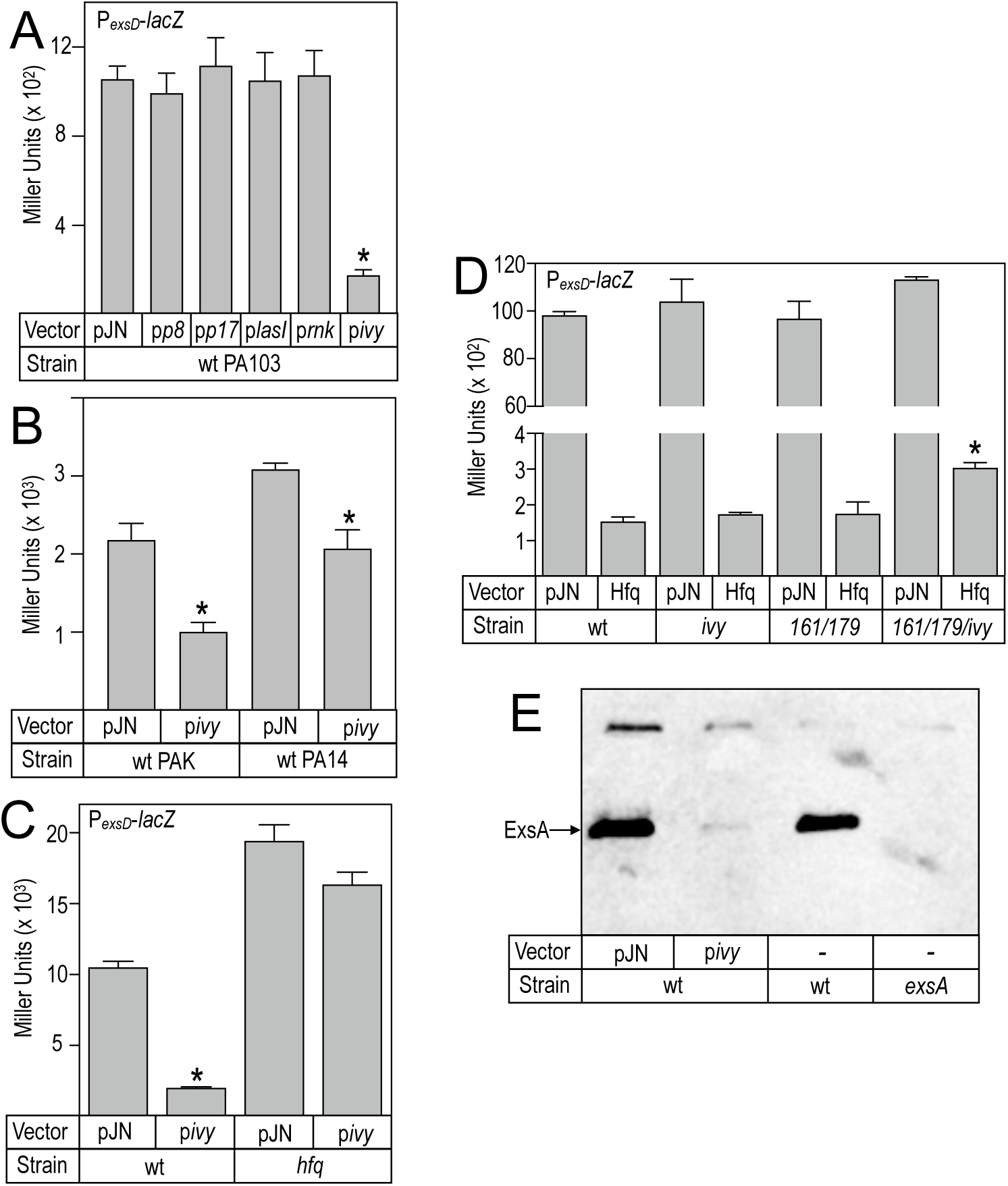
Inhibition of T3SS gene expression by sRNA ivy is Hfq-dependent. **(A)** WT PA103 carrying a P*_exsD_-lacZ* reporter were transformed with either a vector control (pJN105), or expression vectors for sRNAs p8, p17, lasI, rnk, or ivy as indicated. Strains were cultured under T3SS-inducing conditions (+EGTA) supplemented with 0.2% arabinose to induce sRNA expression, and assayed for P*_exsD_-lacZ* reporter activity. **(B)** PAK and PA14 carrying a P*_exsD_-lacZ* reporter were transformed with either a vector control (pJN105) or ivy expression vector, and assayed for reporter activity. Values, with standard error, represent the average of at least three experiments, * denotes *P* < 0.05. **(C)** WT PA103 and an Δ*hfq* mutant carrying a P*_exsD_-lacZ* reporter were transformed with either a vector control (pJN105) or an sRNA ivy expression vector. The resulting strains were cultured under T3SS-inducing conditions with 0.2% arabinose to induce sRNA expression and assayed for P*_exsD_-lacZ* reporter activity. **(D)** WT PA103, Δ*ivy*, Δ0161/179, and Δ*ivy*/0161/179 strains with a P*_exsD_*-lacZ reporter were transformed with either the vector control (pJN105) or an Hfq expression vector. Strains were cultured under T3SS-inducing conditions with 0.2% arabinose to induced Hfq expression and assayed for P*_exsD_*-lacZ activity. **(E)** Overexpression of sRNA ivy inhibits ExsA expression. WT PA103 was transformed with either a vector control (pJN105) or an sRNA ivy expression vector. Strains were cultured under T3SS-inducing conditions with 0.2% arabinose. Wildtype PA103 and an Δ*exsA* strain were used as controls. Cell lysates were immunoblotted for ExsA. Values, with standard error, represent the average of at least three experiments, * denotes *P* < 0.05.

Overexpression of Hfq inhibits T3SS gene expression and presumably acts with sRNAs to control T3SS gene expression (Fig. 5D). Prior to the identification of ivy, at least two sRNAs, 0161 and 179, were known to be Hfq-dependent inhibitors of T3SS gene expression (28, 29). Overexpression of Hfq in a mutant lacking both 0161 and 179, however, still results in complete inhibition of T3SS gene expression (Fig. 5D). To determine whether sRNA ivy, 0161, and 179 collectively account for all Hfq-dependent inhibition of T3SS gene expression, we introduced the Hfq expression plasmid into a mutant lacking all three sRNAs. Even in the triple mutant lacking the ivy, 0161, and 179 sRNAs, Hfq expression still resulted in significant inhibition of P*_exsD_-LacZ* reporter activity. There was, however, a significant but modest decrease in Hfq-dependent inhibition when the single ivy sRNA and double 0161 and 170 sRNA mutants were compared to the triple mutant lacking all three sRNAs (Fig. 5D). These observations indicate that at least three sRNAs inhibit T3SS gene expression in an Hfq-dependent manner (Fig. 5D). Finally, we found that the expression of sRNA ivy in the wt strain reduced ExsA protein levels by more than 90% (Fig. 5E).

### Avenues for defining mechanisms that control T3SS gene expression

The FT-Tn-seq approach significantly expanded the list of genes with potential roles in the control of T3SS gene expression. The remaining challenge will be defining mechanisms that account for control of the T3SS. This will be especially challenging for genes of unknown function. RNA-seq is one tool that could be used to gain mechanistic insight. As a test case we performed RNA-seq on five Tn insertion mutants (*tonB3, gidA,* PA2550, PA3699, and *shaC*) with reduced T3SS gene expression. Each of the Tn mutants and the parental strain were grown under T3SS-inducing conditions and processed for RNA-seq. All five of the Tn mutants demonstrated a significant decrease in T3SS gene expression (Fig. 6 and Dataset-1), further validating data from the primary FT-Tn-seq screen. Genes with reduced expression were fairly restricted in the *tonB3* (29 genes with reduced expression)*, gidA* (43 genes), PA2550 (60 genes), and PA3699 (43 genes) Tn mutants, especially when considering that those totals include ∼30 T3SS gene. The *shaC* Tn mutant had a far more extensive regulon with 443 genes showing reduced expression and 276 genes with increased expression. The hope was that the RNA-seq data would reveal altered expression of genes with known roles in the control of T3SS gene expression. Careful examination of the datasets, however, have revealed no obvious explanation for the reduction in T3SS gene expression. This is not an indictment of the RNA-seq approach, just further demonstration of the complex and diverse mechanisms that regulate T3SS gene expression.

**Figure 6.**
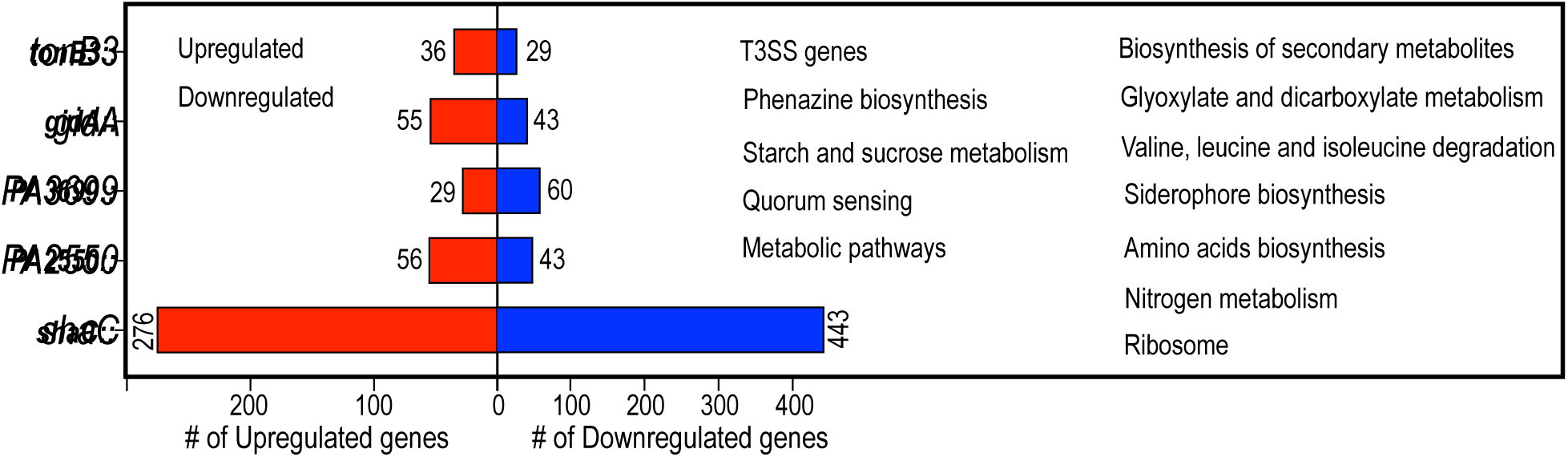
RNA-seq findings for select Tn mutants with defects in T3SS gene expression. RNA-seq analysis confirms that expression of the T3SS regulon is downregulated in select mutant strains identified by the FT-Tn-seq primary screen. Genetic pathways, including the T3SS (green), with altered gene expression in the mutant backgrounds are defined in the legend.

## DISCUSSION

We have been interested in developing a comprehensive screen for genes involved in controlling expression of the *P. aeruginosa* T3SS. In the past we explored screens using the P*_exoS_* promoter to drive expression of either a tetracycline resistance gene (*tetC*) or a gene required for aromatic amino acid biosynthesis (*aroE*) (unpublished). The expectation was that cells capable of T3SS gene expression would become resistant to tetracycline or grow in the absence of aromatic amino acid supplementation, respectively. In both cases, however, the selective pressure imposed for expression of the *tetC* or *aroE* markers resulted in spontaneous P*_exoS_* promoter-up mutations and/or break-through growth of control strains with known defects in T3SS gene expression (eg, an *exsA* mutant). The FT-Tn-seq approach described in this study avoids those issues by using the P*_exoS_* promoter to drive expression of a non-selectable trait (NeonGreen fluorescence). Advantages of this approach include reduced incidence of false positive results and enhanced detection of Tn insertions resulting in smaller effects on T3SS gene expression since there is no threshold level of expression that must be reached to support growth. The P*_exoS_* promoter is highly responsive to inducing condtions with a 150-fold difference in activity when comparing cells grown under non-inducing (-EGTA) to inducing (+EGTA) conditions for T3SS gene expression. This wide range in P*_exoS_* promoter activity contributed to sensitivity by providing a wide window for detection of genes with intermediate effects on T3SS gene expression.

Using a threshold Log_2_FC value of ≤ −1.0 (*p* < 0.05) we identified 73 new genes with potential roles in the control of T3SS gene expression. Confidence in that list is bolstered by several validation studies. First, we examined the 36 T3SS genes encoded continguously on the chromosome. Those genes are required for secretion, regulation, and/or translocation of the effector proteins (8). Two of those genes function as negative regulators of transcription (*exsE* and *exsD*) and are not required for T3SS gene expression (11, 15). Defects in the remaining genes impair transcription or secretory function and should result in reduced T3SS gene expression. Of the 30 T3SS genes (excluding *exsE* and *exsD*) represented in the PA103 library (some genes lacked TA insertion sites), 63% had Log_2_FC values of ≤ −1.0 (*p* < 0.05) (Table 1). Most of the remaining genes also had negative Log_2_FC values but failed to meet statistical significance. The PA103 FT-Tn-seq profile for *exsA*, the primary activator of T3SS transcription, resulted in 7251 sequence reads for the sorted mNG-(dim) population and 752 reads for the mNG+ (bright) profile. This represents a nearly 10:1 signal to noise ratio and indicates that the screening approach was highly sensitive and specific (Table S2). The second validation approach examined 31 non-T3SS genes represented in the PA103 insertion library that were previously known to influence T3SS gene expression (Table 2). From that list, 27 of the 31 genes had negative Log_2_FC values and 21 of the genes met the threshold of ≤ −1.0 (*p* < 0.05) for statistical significance. In the final validation approach we tested 31 *P. aeruginosa* strain PA14 Tn insertion mutants within genes identified by the FT-Tn-seq screen. When tested for T3SS gene expression, 30 of the 31 (97%) insertion mutants demonstrated a significant reduction in expression of a T3SS reporter gene (*p* < 0.05) (Fig. 2). The high validation rate for genes identified in the primary screen using strain PA103 and Tn insertion mutants from PA14 demonstrates the robustness of the approach. Most of those genes are new, with no previously recognized involvement in control of T3SS gene experssion. Potential roles range from direct regulatory functions at the transcriptional and/or posttranscriptional levels to secondary effects that might result from impaired assembly of the type III secretion machine, which is required for maximal T3SS gene expression. Finally the rigor of the study is enhanced through the use of three different *P. aeruginosa* strains (PA103 vs PAK vs PA14). While the magnitude of the effects differed between somewhat between strains for several loci, including components of the cAMP/Vfr and noncanonical regulatory networks, the overall trends were similar in all three strains.

The 70 new genes identified in this study are involved in central metabolism, redox balance, ion homeostasis, cell envelope function, transport, RNA metabolism, and nucleotide signaling. This breadth is consistent with earlier work showing that T3SS expression is sensitive to metabolic state and environmental conditions, rather than being controlled solely by a dedicated virulence regulon (41, 66). Involvement of central metabolism and acetyl-CoA-linked pathways was especially pronounced as summarized in Fig. S2. Carbon flux, redox poise, and energy metabolism may influence bistability and the probability that individual cells transition into a T3SS-On state. This could explain why T3SS expression is heterogeneous even under uniform laboratory inducing conditions: individual cells may differ in metabolic or physiological state, creating permissive or non-permissive states for ExsA-dependent activation.

One of the strongest implicated genes was *shaC*, which encodes a component of a sodium/proton antiporter complex. Transposon insertions in *shaC* were strongly associated with reduced T3SS reporter expression, and targeted deletion of *shaC* reduced maximal T3SS reporter activity in multiple strain backgrounds. The accompanying identification of *shaA* further suggests that Na+/H+ homeostasis, membrane energetics, or intracellular pH may influence T3SS activation. This finding is conceptually consistent with the dependence of T3SS on envelope-associated assembly, proton motive force, and secretion-coupled regulatory feedback. However, the mechanism by which Sha function supports T3SS expression remains unresolved. One possibility is that altered ion homeostasis reduces secretion apparatus activity, thereby preventing secretion of ExsE and limiting liberation of ExsA through the ExsC/ExsD partner-switching cascade. Alternatively, Sha-dependent effects on membrane physiology or cytoplasmic pH may indirectly alter *exsA* expression, translation, or stability. Distinguishing among these models will require direct measurements of secretion, ExsA abundance, intracellular pH, and membrane potential in *sha* mutants.

The screen also linked T3SS expression to c-di-GMP and biofilm-associated regulatory pathways. Mutations affecting *wspF*, a negative regulator of Wsp-dependent c-di-GMP production, reduced T3SS reporter output, and this effect appeared to involve downstream biofilm matrix pathways. These findings fit with a broader regulatory framework in which *P. aeruginosa* coordinates acute virulence traits with surface-associated behaviors and chronic persistence programs. RetS, Gac/Rsm signaling, c-di-GMP, and exopolysaccharide production have all been implicated in balancing T3SS expression, type VI secretion, motility, and biofilm formation (61, 67). Our data extend this concept by showing that FT-Tn-seq can resolve genetic inputs into this balance at the level of single-cell T3SS reporter states. Importantly, the relationship between c-di-GMP and T3SS is unlikely to be strictly binary. Prior work indicates that biofilm-associated traits and T3SS expression are not always mutually exclusive, and the outcome likely depends on strain background, growth condition, c-di-GMP pool localization, and the specific downstream effector pathways engaged (67). Thus, *wspF* mutant phenotypes could be interpreted as evidence that elevated or mislocalized c-di-GMP signaling can constrain T3SS expression under the conditions tested, rather than as a universal rule that c-di-GMP suppresses T3SS in all contexts.

Identification of small RNA-mediated regulation of T3SS expression supports prior studies showing that Hfq and specific sRNAs can repress the T3SS and cAMP/Vfr regulons by reducing ExsA and Vfr synthesis or by modulating the Gac/Rsm pathway (28). Our screen identified candidate sRNA *ivy* as another Hfq-dependent inhibitor of T3SS expression. This expands the known repertoire of post-transcriptional inputs into the T3SS network and suggests that multiple sRNAs may modulate virulence expression in response to distinct physiological cues. Because Hfq-dependent sRNAs can act directly on target mRNAs or indirectly through regulatory cascades, future work should determine whether Ivy represses T3SS by targeting *exsA*, *vfr*, *rsmA*-linked pathways, or another upstream regulator. Mapping Ivy-dependent transcriptome changes and identifying direct RNA-RNA interactions will be important next steps.

This study has several limitations. First, FT-Tn-seq identifies genes whose disruption alters reporter-state distribution, but the screen does not by itself distinguish direct transcriptional regulators from genes that affect growth, secretion apparatus assembly, metabolism, cell envelope integrity, or reporter maturation. Second, genes essential for growth or lacking suitable transposon insertion sites may be missed, and genes with subtle effects on T3SS expression may fall below detection thresholds. Third, the screen was performed under defined *in vitro* T3SS-inducing conditions, which may not capture host-specific signals encountered during infection.

Overall, these findings support a model in which T3SS expression is controlled by an ExsA-centered regulatory module integrated into a broader physiological network. Ion homeostasis, metabolism, c-di-GMP signaling, biofilm matrix pathways, and Hfq-dependent small RNAs all shape the likelihood that individual cells activate T3SS gene expression. This organization may allow *P. aeruginosa* to activate acute virulence in response to local environmental conditions while preserving population heterogeneity. Such heterogeneity could provide a bet-hedging strategy during infection, enabling subsets of cells to deploy cytotoxic T3SS-mediated host interactions while others remain in less inflammatory or more persistent states. By revealing both known and previously unrecognized regulators, FT-Tn-seq provides a generalizable platform for dissecting single-cell heterogeneity in bacterial virulence programs and identifies new regulatory nodes that may be exploited to attenuate *P. aeruginosa* pathogenesis.

## MATERIALS AND METHODS

### Bacterial strains and culture conditions

The bacterial strains used in this study are listed in Table S5. *E. coli* strains were maintained on LB medium with tetracycline (12 μg/ml) or gentamicin (15 μg/ml) as required. *P. aeruginosa* strains were maintained on Vogel-Bonner minimal (VBM) medium supplemented with gentamicin (80 μg/ml), carbenicillin (300 μg/ml), or tetracycline (50 μg/ml) as needed.

### Reporters, plasmids, and strain constructions

Plasmids used in this study are provided in Table S6 and the primers and gene fragments used in their construction are listed in Table S7. PCR products were cloned into plasmids using isothermal assembly. The P*_exoS_*-mNeonGreen (P*_exoS_*-mNG) reporter was constructed by combining the sequences of the exoS promoter, ribosomal binding sites, and mNeonGreen using a gBlock approach. The gBlock gene (Table S7) was cloned into pUC18-mini-Tn7T-GM-LacZ10 (Addgene #65026) using isothermal assembly. The resulting Tn7 P*_exoS_*-mNG plasmid was introduced into *P. aeruginosa* by electroporation using the pTNS2 helper plasmid (Addgene #64968) (68) and integrated intoi Tn7 chromosome attachment site. Allelic exchange vectors for deleting *shaC*, PA2550, PA3699, PA5435, *wspF*, *wspR^D16N^* variant, and ivy were generated by PCR using primers summarized in Table S7. PCR products were cloned by isothermal assembly into pEXG2Tc. The resulting constructs were mobilized from *E. coli* SM10 into *P. aeruginosa* via conjugation. Merodiploids were selected on VBM agar with tetracycline (50 μg/ml) and resolved by counterselection using the *sacB* marker on YT medium with 10% sucrose.

### FT-Tn-seq approach

*P. aeruginosa* carrying the P_exoS_-mNG reporter was analyzed and sorted using a Becton Dickinson Aria Fusion flow cytometer with a 70 μm nozzle, 70 PSI pressure, and the GFP fluorochrome combination (525/50 BP). *P. aeruginosa* cells were cultured overnight in LB broth and back-diluted the next day to OD_600_ 0.1 in LB broth. To induce mNG T3SS reporter activity, the media were supplemented with either 2 mM (PA103) or 10 mM (PAK) EGTA. When cultures reached an OD_600_ of 0.6-0.8, 1 ml of cells were collected by centrifutation (5000 x *g* for 5 min), washed with 1 ml PBS, diluted 1:5 in PBS, held on ice, and directly analyzed with flow cytometry. For the first gating control, *P. aeruginosa* lacking the mNG reporter was grown in LB broth supplemented with EGTA. For the second gating control, *P. aeruginosa* strain harboring the mNG reporter was grown in LB broth lacking EGTA. Flow cytometry was used to monitor the mNG reporter at all steps.

*P. aeruginosa* strains harboring the mNG reporters were mutagenized as previously reported (69, 70). The Mariner carrying pBT20 vector (46) was mobilized via conjugation from *E*. *coli* SM10 λpir to the recipients PA103 Δ*exoUT* P_exoS_-mNG or PAK Δ*exoSTY* P_exoS_-mNG strains. The conjugation reactions were plated on *Pseudomonas* isolation agar and cells from 75 or 150 plates were pooled to create the PA103 and PAK mutant libraries, respectively.

For flow cytometry analyses, the PA103 transposon insertion library was thawed on ice, and approximately 3 x 10^9^ cells were inoculated into 50 ml LB broth supplemented with 2 mM EGTA. A parallel culture without EGTA served as an input control. The cultures were incubated in 250 ml Erlenmeyer flasks at 37°C and 200 RPM until the OD_600_ reached 0.6. The PAK insertion library was inoculated in 500 ml LB medium supplemented with 200 mM NaCl, 10 mM MgCl_2_, and 10 mM EGTA in a 2.8 L flask and grown overnight. The following morning, the culture was diluted 1:50 in LB and grown to an OD_600_ of 0.9 in a 50 ml culture. Cells were sorted in PBS and mixed with an equal volume of 2X LB both. The shorted dim and bright cells were separately cultured to an OD_600_ of 0.5. Cells (2 ml) were then pelleted and stored at −20°C for gDNA extraction. DNA libraries were prepared for Illumina sequencing as previously reported (69, 70).

### FT-Tn-seq data analyses

Sequencing data from the libraries was demultiplexed based on barcodes and a modified TPP software tool was used to remove sequences upstream of the genomic junctions, trim the 3’ c-tails if present, and map the sequences to the reference PA103 genome using BWA 0.7.17 (71, 72). To facilitate our analyses we completed the genome sequence of the PA103 strain using long-read and short-read sequencing. The genome of *P. aeruginosa* strain PA103 was sequenced using Illumina short-read sequencing as described previously (73) and long-read sequencing with Oxford Nanopore technology. Sequencing adaptors and low-quality reads were removed using Cutadapt (74). The complete circular genome of PA103 was assembled using Trycycler (75) and is available in the NCBI database under accession number CP136909. Open reading frames and small non-coding RNAs were annotated with Prokka and Infernal (63, 76). The complete genome of PA103 spans 6,791,681 bp and encompasses 6,198 genes, 65 tRNAs, 12 rRNAs, and 87 ncRNAs.

We used the completed PA103 genome as the reference for PA103, and the *P. aeruginosa* PAK genome (NZ_LR657304) as the reference for PAK (77). TRANSIT software was used to compare number of reads in dim and bright mutant and to calculate Log_2_FC for the genes and sRNAs (71).

### β-Galactosidase assays and immunoblots

*P. aeruginosa* strains were cultured overnight at 37°C in LB with 80 μg/ml gentamicin as required. The next day cells were diluted to an OD_600_ of 0.1 in trypticase soy broth (TSB) supplemented with 100 mM monosodium glutamate,1% glycerol, 80 μg/ml gentamicin, and 2 mM ethylene glycol tetraacetic acid (EGTA). Arabinose/rhamnose was added to induce the gene or sRNA of interest as required. Cells were then grown to an OD_600_ of 1.0 and β-galactosidase activity was assayed with either ortho-nitrophenyl-galactopyranoside (ONPG) or chlorophenol red-β-D-galactopyranoside (CPRG) substrates as previously described (18). Immunoblots using rabbit immune serum to ExsA was performed as previously described (11). P*_cdrA_*-GFP fluorescent reporter was measured as previously reported (60).

### RNA Sequencing

*P. aeruginosa* PA14 wt and Tn insertion mutants in *shaC*, PA2550, PA3699, *gidA*, or *tonB3* were grown under T3SS-inducing conditions with 2 mM EGTA until reaching an OD_600_ of 1.0. RNA was extracted using the SPLIT RNA extraction kit. Ribosomal RNA was depleted using the RiboCop rRNA depletion kit and libraries were prepared using the CORALL RNA-Seq library prep kit (Lexogen). Libraries were sequenced on the Illumina HiSeq X platform. Sequencing read adaptors were trimmed, and low-quality reads were removed using Cutadapt (74). RNA-seq analyses were conducted on the BV-BRC system using the Tuxedo strategy (78).

## Statistical analysis

Student two-tailed unpaired tests were performed using Prism 10 GraphPad.

## Data availability

Tn-seq sequencing reads are available on NCBI Sequence Read Archive under BioProjects PRJNA849614 for PA103 and PRJNA849930 for PAK.

## ACKNOWLEDGMENTS

This work was supported by National Institutes of Health grant number R01 AI055042-15 to T.L.Y. Special thanks to flow cytometry facility and high-performance computing at the University of Iowa. Thanks to Matthew Radey for assembling the complete genome of PA103.

---

**Table 1.** Genes with known roles in T3SS gene expression identified by FT-Tn-seq in *P. aeruginosa* strains PA103 and PAK.

**Table 2.** FT-Tn-seq identified new genes that positively regulate the expression of genes required for T3SS function in *P. aeruginosa* strains PA103 and PAK.

**Table S1.** Number of sequencing reads and unique insertion mutants utilized in FT-Tn-seq.

**Table S2.** Comprehensive list of genes identified by FT-Tn-seq in *P. aeruginosa* strain PA103 that positively regulate T3SS expression.

**Table S3.** Comprehensive list of genes identified by FT-Tn-seq in *P. aeruginosa* strain PAK that positively regulate T3SS expression.

**Table S4.** List of sRNAs identified by FT-Tn-seq in *P. aeruginosa* strain PA103 that regulate T3SS expression, with sRNA sequences listed.

**Table S5.** Bacterial strains used in this study.

**Table S6.** Plasmids used in this study.

**Table S7.** gBlocks and primers used in this study.

**Figure S1. Validation of P*_exoS_*-mNeonGreen reporter.** WT *P. aeruginosa* strains PA103 **(A** and **B)** and PAK **(D** and **E)** carrying the P*_exoS_*-mNG reporter were cultured under non-inducing (-EGTA) and inducing (+EGTA) conditions for T3SS gene expression. The percentage of cells expressing mNG+ (T3SS-On) was quantified using flow cytometry. (C and F) The reporter intensity remained stable when the same experiment was performed using the effectorless PA103 Δ*exoUT* **(C)** and PAK Δ*exoSTY* **(F)** mutants. **(G)** Cells from the PAK strain passaged twice in +EGTA maximized induction of the P*_exoS_*-mNG reporter. Values, with standard error, represent the average of at least three experiments, * denotes *P* < 0.05.

**Figure S2. Acetyl Co-A in central metabolic pathways modulates T3SS expression.** FT-Tn-seq identified multiple genes involved in acetyl Co-A generation within central metabolism that regulate T3SS expression. Genes highlighted in red were identified by FT-Tn-seq, with those in bold exhibiting Log_2_FC with *P* < 0.05.

